# N-acylethanolamine acid amide hydrolase is a novel target for drugs against SARS-CoV-2 and Zika virus

**DOI:** 10.1101/2022.02.08.479661

**Authors:** Michele Lai, Veronica La Rocca, Rachele Amato, Elena Iacono, Carolina Filipponi, Elisa Catelli, Lucia Bogani, Rossella Fonnesu, Giulia Lottini, Alessandro De Carli, Alessandro Mengozzi, Stefano Masi, Paola Quaranta, Pietro Giorgio Spezia, Giulia Freer, Paola Lenzi, Francesco Fornai, Daniele Piomelli, Mauro Pistello

**Author notes:** Authors provided equal contribution (Michele Lai, Veronica La Rocca, Daniele Piomelli, Mauro Pistello). Corresponding author:* Michele Lai, Centro Retrovirus; Dipartimento di Ricerca Traslazionale e delle Nuove Tecnologie in Medicina e Chirurgia, Università di Pisa, SS 12 Abetone e Brennero, 2, Pisa, I-56127, Italy.

## Abstract

Several compounds have been tested against SARS-CoV-2; at present, COVID-19 treatments decrease the deleterious inflammatory response and acute lung injury. However, the best therapeutic response would be expected by combining anti-inflammatory properties, while concomitantly blocking viral replication. These combined effects should drastically reduce both infection rate and severe complications induced by novel SARS-CoV-2 variants. Therefore, we explored the antiviral potency of a class of anti-inflammatory compounds that inhibit the N-Acylethanolamine acid amidase (NAAA). This enzyme catalyzes the hydrolysis of palmitoylethanolamide (PEA), a bioactive lipid that mediates anti-inflammatory and analgesic activity through the activation of peroxisome proliferator receptor-α (PPAR-α). Similarly, this pathway is likely to be a significant target to impede viral replication since PPAR-α activation leads to dismantling of lipid droplets, where viral replication of Flaviviruses and Coronaviruses occurs.

Here, we show that either genetic or pharmacological inhibition of the NAAA enzyme leads to five-fold reduction in the replication of both SARS-CoV-2 and ZIKV in various cell lines. Once NAAA enzyme is blocked, both ZIKV and SARS CoV-2 replication decrease, which parallels a sudden five-fold decrease in virion release. These effects induced by NAAA inhibition occurs concomitantly with stimulation of autophagy during infection. Remarkably, parallel antiviral and anti-inflammatory effects of NAAA antagonism were confirmed in *ex-vivo* experiments, within SARS-CoV-2 infected human PBMC cells, in which both viral genomes and TNF-α production drop by ~60%. It is known that macrophages contribute to viral spread, excessive inflammation and macrophage activation syndrome that NAAA inhibitors might prevent, reducing the macrophage-induced acute respiratory distress syndrome and subsequent death of COVID-19 patients.

## Introduction

SARS CoV-2 pandemic is a major emergency for public health worldwide. So far, the virus caused hundreds of thousands of deaths out of 260 million infection cases (WHO Dashboard). Since 2019, nine vaccines have been approved for use in humans(1). Two of the vaccines currently in use worldwide, BNT162b2 (manufactured by Pfizer) and mRNA-1273 (manufactured by Moderna), are based on mRNA encoding the spike protein derived from SARS-CoV-2 isolated early in the epidemic from Wuhan, China. Both vaccines demonstrated >94% efficacy at preventing coronavirus disease 2019 (COVID-19)(2, 3). However, recent emergence of novel SARS-CoV-2 variants has raised significant concern about the geographic and temporal efficacy of these interventions.

Even though several compounds have been tested against SARS CoV-2, current COVID-19 treatments aim at decreasing the severe inflammatory response and acute lung injury in patients. Unfortunately, blocking the immune system during viral infections might end up being advantageous on the viral side and increase the viral spread. In fact, after inflammation, immune cells also generate beneficial effects to self-limit inflammation and promote tissure recovery. In addition such a mechanism does not occlude the process of viral replication. Thus, additional effects, which may block viral replication are required to encompass all facets of an effective cure. Thus, we are in need of drugs, which may combine the mitigation of a drastic inflammatory process along with occluding viral replication independently by strain specificity. This is expected to reduce disease severity and confine effectively the spreading of novel variants.

All mammalian cells express a variety of signaling molecules to communicate with their surroundings and boost the inflammatory response. In addition to cytokines, peptides, and small molecules, several signaling lipids are also involved in the regulation of inflammation. Lipid messengers involved in reconstructive actions include lipoxins, resolvins, and the N-acylethanolamines (NAE), such as arachidonoylethanolamide (anandamide) and palmitoylethanolamide (PEA)(4). Anandamide is the best characterized NAE, together with its related compound 2-arachidonoylglycerol, and is known to activate cannabinoid (CB) 1 and CB2 receptors (5). On the contrary, PEA, a well-recognized analgesic, anti-inflammatory, and neuroprotective mediator, does not activate CB receptors(6). PEA exerts its anti-inflammatory action through activation of peroxisome proliferator-activated receptor-α (PPAR-α)(7), a ligand-activated transcription factor. Significantly. marked decrease in PEA content has been described in inflammatory exudates of endotoxin-exposed mice(8).

N-Acylethanolamine acid amidase (NAAA) is an N-terminal cysteine hydrolase primarily found in the endosomal-lysosomal compartment of innate and adaptive immune cells(9). NAAA catalyzes the hydrolytic cleavage of PEA. Considering the analgesic and anti-inflammatory actions of PEA, multiple NAAA inhibitors have been developed as anti-inflammatory drugs over the last decade(10). These compounds exhibit beneficial effects in a range of rodent models of human diseases, including inflammation(11), neuropathic pain(12), chronic pain(12), and lung inflammation(13). NAAA is part of the N-terminal nucleophile (NTN) hydrolase superfamily, a diverse group of amidases that carry out catalysis via their N-terminal cysteine, serine, or threonine residues(9). NAAA is produced as a precursor and undergoes activation by self-cleavage of an internal peptide bond. This results in the formation of the mature enzyme composed of an α- and a β-subunit that remain associated. Due to their lipidic nature, N-acylethanolamines are transported via carrier proteins, membrane vesicles, or lipid droplets. Although NAAA is a soluble protein, the enzyme disrupts vesicular membranes to reach PEA thanks to its α3 and α6 helices, bearing almost exclusively hydrophobic and cationic side chains, allowing NAAA to act as a monotopic membrane protein.

NAAA inhibitors exert a profound anti-inflammatory activity, comprising the reduction of IL-6 and TNF-α secretion by macrophages(14). In contrast, their role during viral infections is still poorly characterized. Although NAAA inhibitors may interfere with the replication of SARS-CoV-2 and flaviviruses due to the same mechanism that NAAA inhibitors exert to control inflammation(15), alternative mechanisms should be considered. These include the biochemical events triggered by PPAR-α activation. Indeed, PPAR-α generates a cascade of signalling that leads to the disruption of fatty acid droplets by the activation of β-oxidation within mitochondria and peroxisomes, and concomitant stimulation of omega-oxidation in microsomes(16). These lipid droplets are essential for the replication of flaviviruses(17) Indeed, once flaviviruses are internalized, a combination of membrane-associated viral proteins and host proteins cooperate to create clusters of invaginated vesicles, called membranous web, in which host lipids shield dsRNA from cytoplasmic sensors(17). As recently suggested, the membranous web is also important for SARS CoV-2 replication(18).

In this work, we assess whether NAAA inhibitors affect the replication of viruses that uses lipid droplets to boost their replication, such as Zika (ZIKV) and SARS CoV-2 and whether this is due their ability to dismantle the lipid compartments, which are supposed to sustain the membranous web. Moreover, we questioned whether activation of PPAR-α might sustain autophagy progression, thus contributing to the elimination of viral proteins, thus posing a second line of resistance against virus spreading.

## Results

### NAAA expression increases after ZIKV infection

To probe the hypothesis that NAAA is exploited during flavivirus infection, we measured NAAA mRNA before and after ZIKV infection in A549 and Huh-7cells, two well-established cellular models for virological studies. As shown in Figure 1a, we infected both A549 and Huh-7 with ZIKV at 1 multiplicity of infection (MOI) for 1 h. As a control, we transfected cells with polyI:C, a synthetic double-stranded RNA that mimics viral intermediate RNA molecules and activates the intrinsic cellular response(19). Then, we collected total RNA at 24 and 48h post-infection. As shown in Figure 1b, both A549 and Huh-7 cells increase NAAA expression by ~2 and 3 folds at 48h, respectively, whereas little difference was found at 24h, but not in polyI:C-transfected ones. To confirm NAAA up-regulation at a protein level, we probed ZIKV-infected A549 cells with both anti-NAAA and anti-ZIKV antibodies by high content confocal immunofluorescence. As shown in Figure 1c-d, increased NAAA expression could only be detected in ZIKV-infected cells.

**Figure 1.**
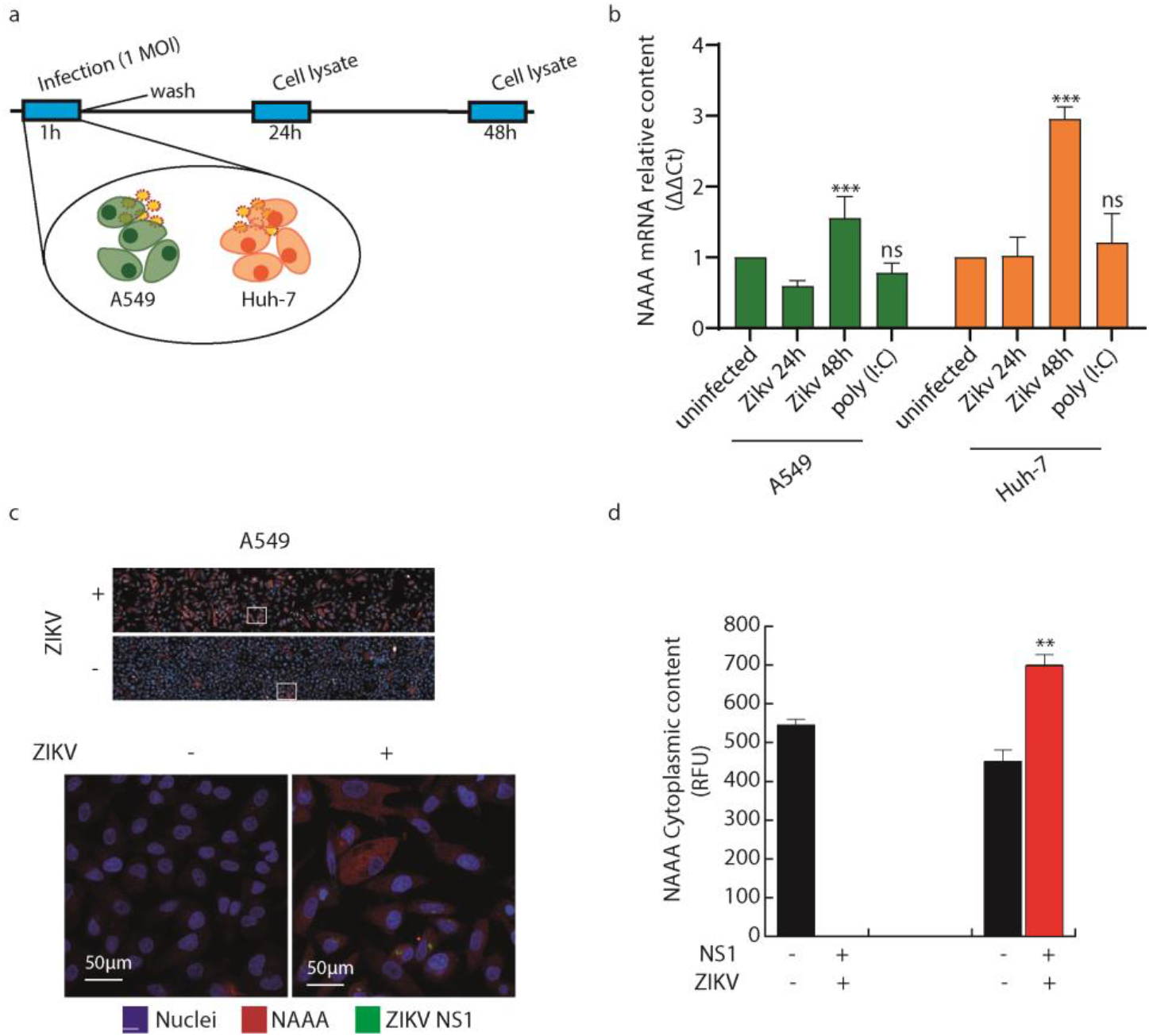
NAAA expression increases after ZIKV infection. a) Schematic illustration of the experimental workflow. Briefly, A549 and Huh-7 cells were infected with 1 MOI of ZIKV and, 24 and 48h later, NAAA mRNA levels were measured in cell lysates. b) NAAA mRNA content, measured by qRT-PCR, in ZIKV infected lysates at 24 and 48h after infection. Data are expressed as mean ± SD, N=3, normalized with β-actin, and analyzed by One-way Anova (*** p<0.001, ** p<0.01). c) Immunofluorescence staining performed by high-content confocal microscopy of A549 cells infected or not with 1 MOI of ZIKV, 48h post-infection. Upper panel: an overview of 30 fields; lower panel 63× magnification of the white squares in upper panels. Nuclei were stained with DAPI (blue), with ZIKV NS1 Alexa 488 (green), and NAAA with Alexa 688 (red). d) Statistical analysis of high content screening shown in c. Data are expressed as mean RFU ± SD N=3, and analyzed by One-way Anova (** p<0.01).

### ZIKV NS1 co-localizes with NAAA and autophagosomes

Due to their lipid characteristics, N-acylethanolamines are not likely to be freely available in the cytoplasm; rather, they are transported *via* carrier proteins, membrane vesicles, or lipid droplets(20). Consequently, also their degrading enzymes must be able to access these compartments within these hydrophobic settings. Several lines of evidence show that NAAA localizes in lysosomes(11). We hypothesized that this enzyme might also be present in other vesicular compartments, such as endosomes and autophagosomes. Considering that ZIKV entry occurs exclusively through endocytosis(21) we monitored the localization of ZIKV NS1 protein and NAAA, using high-content confocal microscopy. As shown in Figure 2a, we infected A549 cells with ZIKV at 5 and 10 MOI, to maximize the number of infected cells. Analyses performed 24h after infection revealed that ZIKV NS1 co-localizes with NAAA in ~30% of cytoplasmic vesicles (Figure 2b-d). To determine the nature of these vesicles, we took advantage of a fluorescence-based autophagy assay (LC3-GFP RFP assay), where the autophagic marker LC3, expressed as a fusion protein with two fluorochromes, allows to distinguish autophagosomes (GFP+/RFP+ vesicles) from autolysosomes (GFP-/RFP+ vesicles) in transfected cells(22, 23). In this assay, LC3-GFP fluorescence is quenched when pH drops due to lysosome-autophagosome fusion. This assay, schematically illustrated in Figure 2e, shows that NAAA localizes in both autophagosomes (~11% and 14% for A549 and Huh-7, respectively) and autolysosomes (~11% and 7% for A549 and Huh-7, respectively).

**Figure 2.**
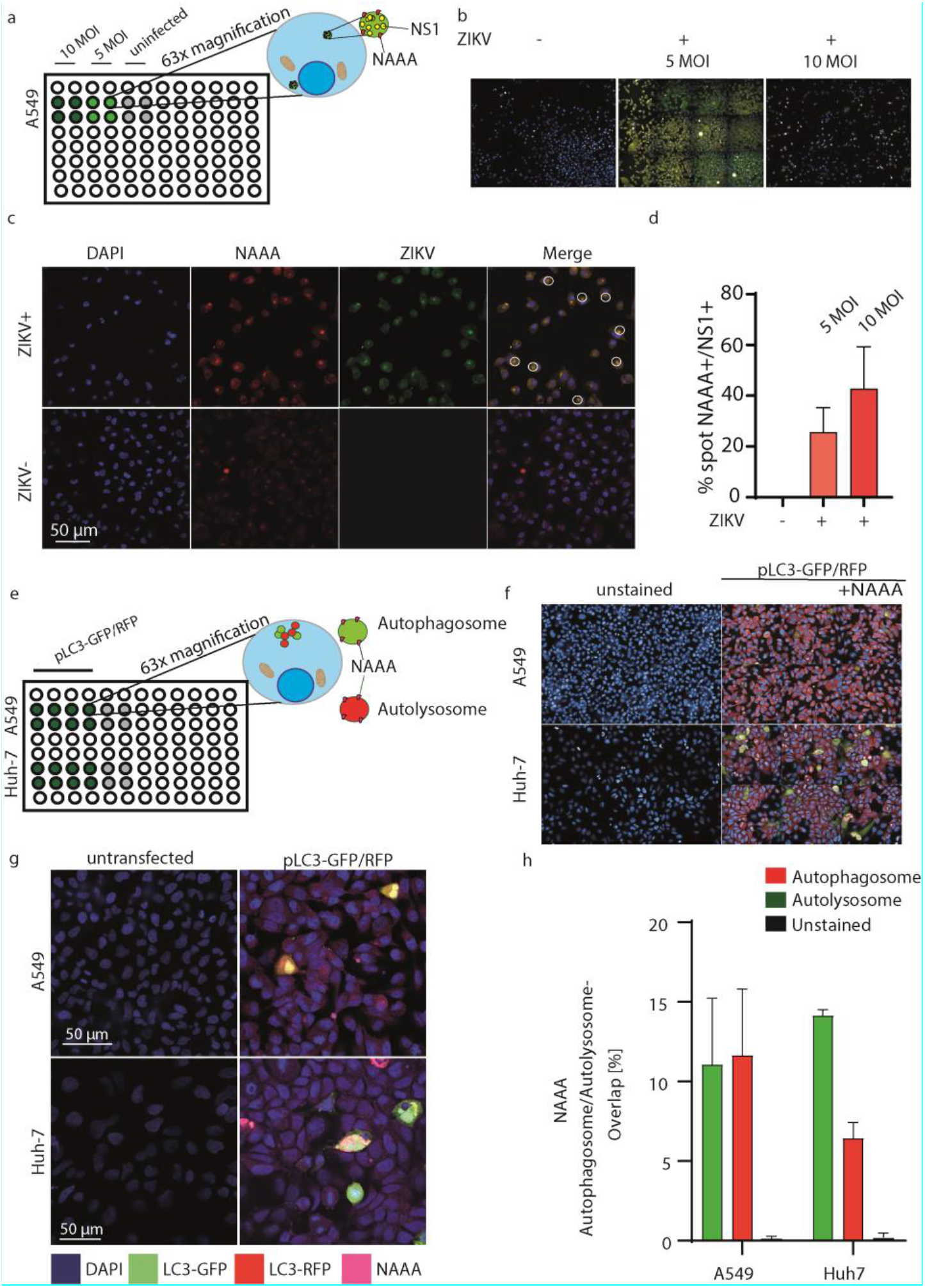
ZIKV NS1 co-localizes with NAAA in vesicles. **a)** Schematic high content confocal screening performed on A549, and Huh-7 cells infected or not with 5 or 10 MOI of ZIKV. Cells were fixed 24h after infection, stained for NS1 and NAAA, and probed for spot co-localization. Twenty-five fields were analyzed per well using a 63× water objective. **b)** Overview of 12 fields from high-content confocal screening. Left: uninfected cells, center: 5 MOI ZIKV, right: 10 MOI ZIKV. **c)** Representative images are taken from the acquisition shown in b. NAAA is marked in red, nuclei in blue, and ZIKV NS1 in green. **d)** Statistical analysis of NS1-NAAA spot overlap, data are expressed as a mean count of overlapping spots ± SD, N=4, alpha= 0.5. **e)** Schematic high content confocal screening performed on uninfected A549 and Huh-7 cells. Cells were transfected with pLC3-GFP/RFP, then fixed 24h after. NAAA co-localization with autophagosomes (GFP+/RFP+ spots, yellow) and autolysosomes (GFP-/RFP+ spots, red) was performed using the following building block: find nuclei > find cytoplasm > Calculate intensity properties (GFP, RFP, NAAA (Alexa 647) > find spots (GFP+RFP, GFP-/RFP+) > calculate position properties (% overlap GFP+/RFP+ and NAAA, GFP-/RFP+ and NAAA). Twenty-five fields were analyzed per well using a 63× water objective. f) Overview of the acquisition described in e. g) Representative images of the acquisition described in e-f. h) Statistical analysis performed on % overlap of NAAA with autophagosomes (GFP+/RFP+) and autolysosomes (GFP-/RFP+) using One-way Anova. Data are expressed as mean ± SD, N=3, alpha= 0.5.

### Ablation of NAAA interferes with ZIKV infection

We used CRISPR/Cas9 genome editing to generate NAAA -/- A549 and Huh-7 cell lines bearing frameshift mutations in both NAAA alleles. Production of NAAA-/- cell clones is described in Supplementary Information.

To evaluate the effect of NAAA ablation on ZIKV replication, we infected A549 NAAA-/- clones at a low MOI (0.1 MOI) of ZIKV, to avoid excessive cellular stress. It is well known that ZIKV enters cells through the endocytic pathway only (24) (Figure 3a), making it highly likely to co-localize with NAAA during entry.

**Figure 3.**
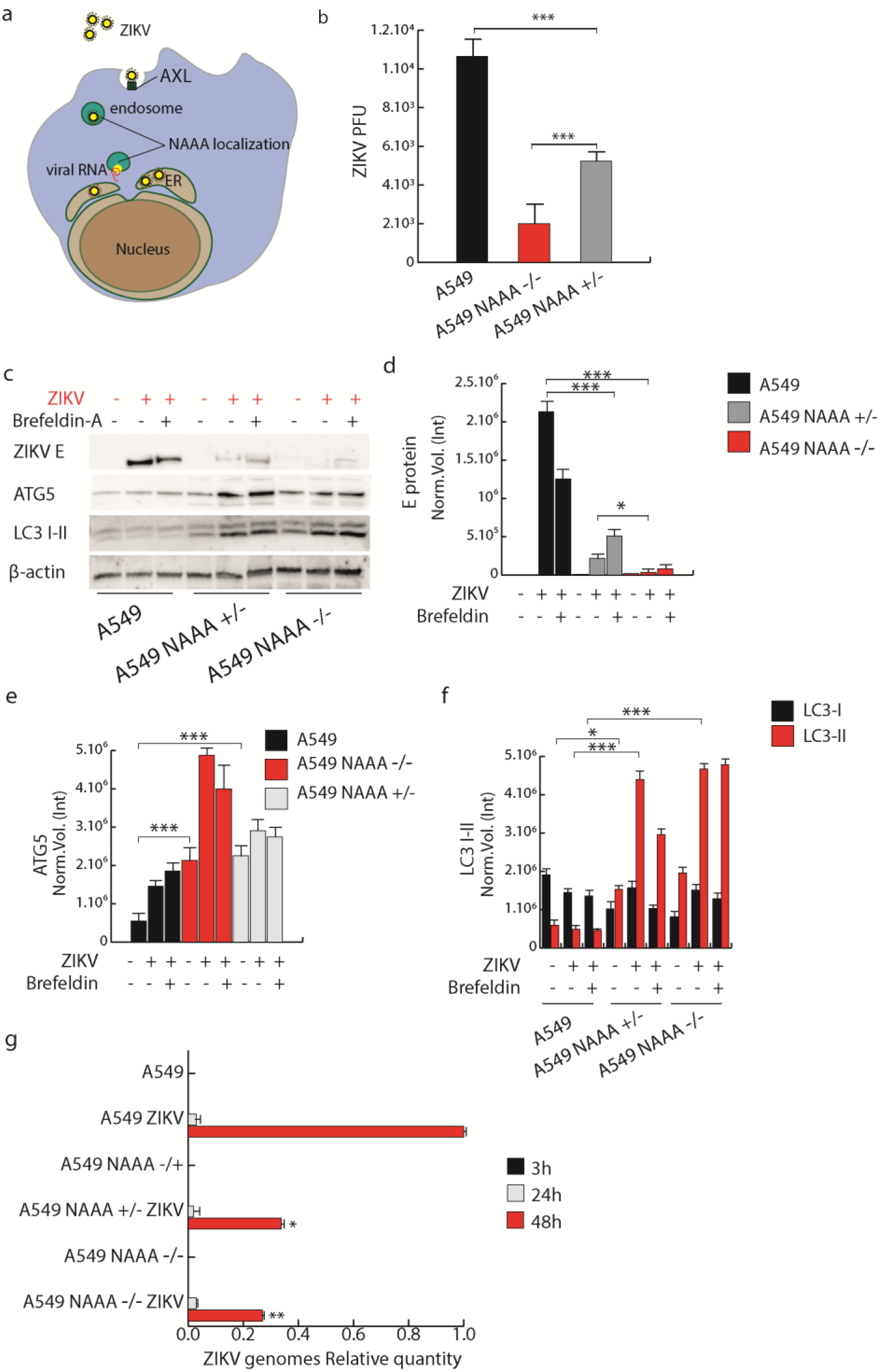
NAAA ablation reduces ZIKV replication. **a)** Schematic illustration of ZIKV entry through the endocytic pathway. **b)** Plaque assay performed on A549 and A549 NAAA-/- cells infected with 0.1 MOI of ZIKV. **c-f)** Western blot performed on A549, A549 NAAA+/- and A549 NAAA-/- using anti-ZIKV E, ATG5 and LC3 I-II. Statistical analysis (d-f) was performed using One-Way Anova (p<0.01 *; p<0.001 **; p<0.0001 ***, α=0.05). Data are expressed as mean ± SD, N=4. **g)** qRT-PCR quantification of ZIKV viral genomes performed on supernatants of A549, A549 NAAA+/- and A549 NAAA-/-, taken at different time-points (3h, 24h, 48h). Statistical analysis was performed using One-Way Anova (p<0.01 *; p<0.001 **; p<0.0001 ***, α=0.05). Data are expressed as mean ± SD, N=3.

Figure 3b shows that A549 NAAA-/- cells exhibit a 5-fold reduction in the number of ZIKV plaque-forming units (PFU) compared to WT controls. Interestingly, we observed a moderate reduction of ZIKV PFUs also in the NAAA heterozygotic A549 clone, where NAAA content is significantly reduced but not abolished. Western Blot analysis confirmed a decrease in viral content since a 20-fold reduction in ZIKV E intracellular protein content was measured in both A549 NAAA -/- and NAAA +/-, compared to controls (Fig 3c-d). Diminution of ZIKV E protein was enhanced by the addition of Brefeldin-A, a drug that inhibits protein transport from the endoplasmic reticulum to the Golgi complex, leading to intracellular ZIKV E accumulation (Fig 3c-d).

During viral shutoff, ZIKV uses autophagosomes to continue its assembly(25), however, the virus impedes the progression of the autophagy flux to prevent autophagy-mediated virion disruption, This effect is achieved through the inhibition of autophagosome fusion with lysosomes, thus impairing the eventual autophagy flux (26). To test whether NAAA depletion might interfere with the ZIKV-induced blockade of autophagy flux, we measured autophagy in NAAA -/- clones during ZIKV infection. Accordingly, as shown in Figure 3e-f, A549 NAAA-/- contains twice the level of the autophagy initiator ATG5 protein compared to controls (Fig 3c and e). The induction of autophagy was confirmed by measuring increased levels of LC3-II, which is the reference marker for autophagosome maturation(27). In line with the previous result, LC3-II increases 3-fold in A549 NAAA-/- cells compared to controls, also during ZIKV infection (Fig 3f). Finally, we measured ZIKV genomes in supernatants of each cell type by qRT-PCR at 3h, 24h, and 48h post-infection. To this aim, we added the 3-h time point that allows subtracting the virions used for the infection itself from the count of the virions produced. As shown in Figure 3g, A549 NAAA-/- cells released a 5-time lower number of particles in the supernatants, compared to WT counterparts.

### NAAA inhibitor ARN077 interferes with ZIKV replication

Given the negative effect of NAAA reduction on ZIKV replication, we decided to test the antiviral activity ARN077, a highly potent NAAA inhibitor(12) on A549 cells. The drug was administered 15 minutes before ZIKV infection at the concentration of 0.3, 1, 3, and 10 μM. Western blot analysis (Figure 4a-b) shows dose-dependent ZIKV E protein reduction starting from 0.3 μM ARN077 treatment. Concomitantly with ZIKV E protein reduction, we observed an increase in the autophagy markers LC3-II, Atg5 and p62 when ARN077 is administered (Fig. 4c-e, respectively). Finally, we quantified ZIKV genomes in the supernatants of A549 treated with increasing doses of ARN077 by qRT-PCR (Figure 4f). The analysis revealed a dose-dependent decrease of ZIKV genomes after ARN077 treatment. These results are consistent with the ones obtained with NAAA-/- cells

**Figure 4.**
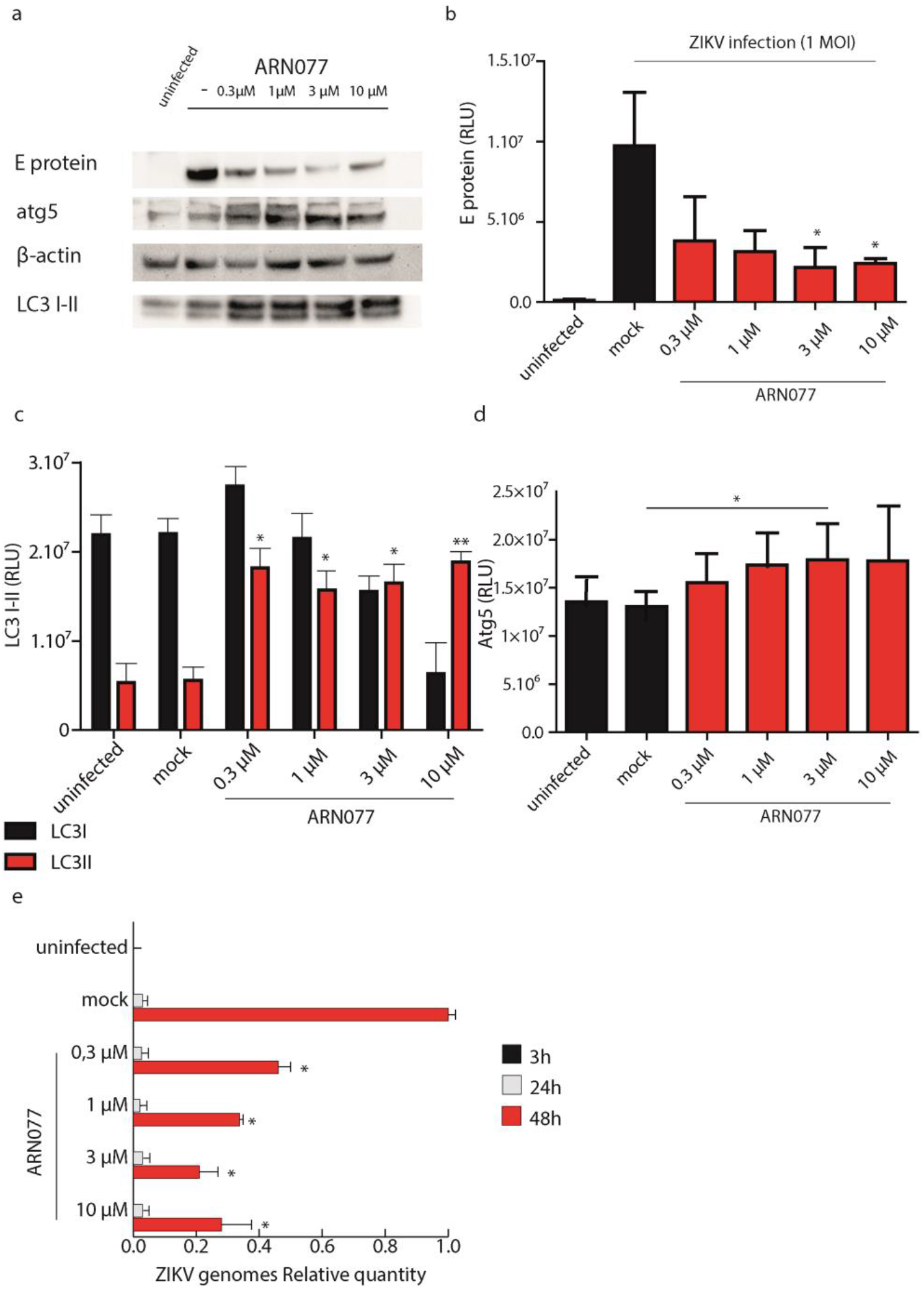
NAAA inhibitor ARN077 decreases ZIKV replication. **a-d)** Western blot performed on A549 cell lysates treated with increasing doses of ARN077 15 min before ZIKV infection, using anti-ZIKV E, ATG5, and LC3 I-II. Statistical analysis was performed using One-Way Anova (p<0.01 *; p<0.001 **; p<0.0001 ***, α=0.05). Data are expressed as mean ± SD, N=4. **e)** qRT-PCR quantification of ZIKV viral genomes performed on supernatants of A549 treated with increasing doses of ARN077 15 min before infection. Supernatants were taken at different points (3h, 24h, 48h). Statistical analysis was performed using One-Way Anova (p<0.01 *; p<0.001 **; p<0.0001 ***, α=0.05). Data are expressed as mean ± SD, N=5.

### NAAA inhibition does not affect the replication of a naked virus

To better elucidate the antiviral specificity of NAAA inhibition, we tested ARN077 on A549 cells infected with 0.1 MOI of Coxsackievirus B5 (COXB5), a small non-enveloped RNA virus belonging to *Picornaviridae*. As shown in Figure 5a, we opted for COXB5 because it replicates in the cytoplasm without strictly depend on double membrane vesicles and autophagy-derived membranes such as ZIKV and SARS-CoV-2. Indeed, while SARS-CoV-2 and ZIKV block the autophagic flux, COXB5 enhances autophagy to promote its replication. As shown by Figure 5b-c, although NAAA inhibition slightly reduced the amount of intracellular coat protein, it was inefficient at reducing viral release, as observed by qR-T PCR performed on cell supernatants (Figure 5d).

**Figure 5.**
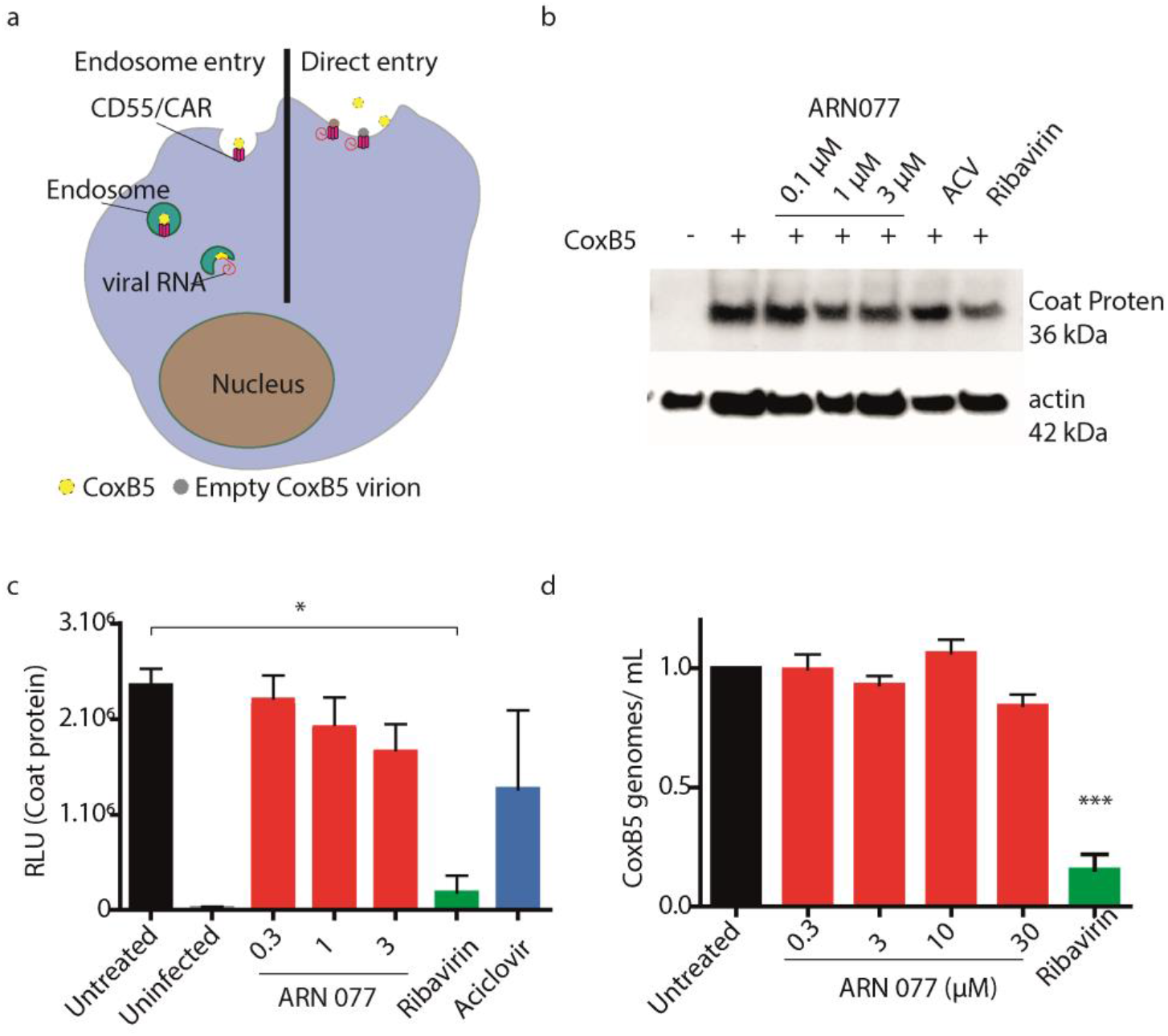
NAAA inhibition does not interfere with COXB5 replication. **a.** Schematic illustration of COXB5 entry mechanism. Briefly, COXB5 virions use both endosome-mediated entry and direct plasma membrane genome release. **b.** Western blot detection of COXB5 coat protein performed on cell lysates taken 48h after infection. **c.** Statistical analysis of COXB5 coat protein, normalized on actin. Data are expressed as mean ± SD, N=3 and analyzed with One-way Anova (* p<0.05). **d.** qRT-PCR quantification of COXB5 genomes performed on cell supernatants, 48h after infection. Data are expressed as mean ± SD, N=3 and analyzed with One-way Anova (*** p<0.001).

### NAAA ablation decreases SARS CoV-2 replication

Since ZIKV share similar entry mechanisms and similar involvement of lipid droplets of SARS-CoV-2, we assess if NAAA ablation might exert the same antiviral activity against SARS-CoV-2 (Figure 5a). To this aim, we infected Huh-7 NAA-/- cells with 1 MOI of SARS-CoV2, Wuhan strain. Cells were exposed to infection for 1h, then washed and kept on fresh media for 48h. Media was collected at 24h and 48h to follow viral replication.

Figure 5b shows that Huh-7 NAAA-/- cells decrease the release of SARS-CoV-2 virions in the supernatant by 5-fold compared to WT cells. This was accompanied by a 3-fold reduction in viral N protein content in cell lysates (Figure 5c-d). Concomitantly, in Huh-7 NAAA-/- cells autophagy activation occurred, which was measured by LC3-II and p62 content, which parallels what described in NAAA-/- cells during ZIKV infection (Figure 5 b-d-e).

**Figure 5.**
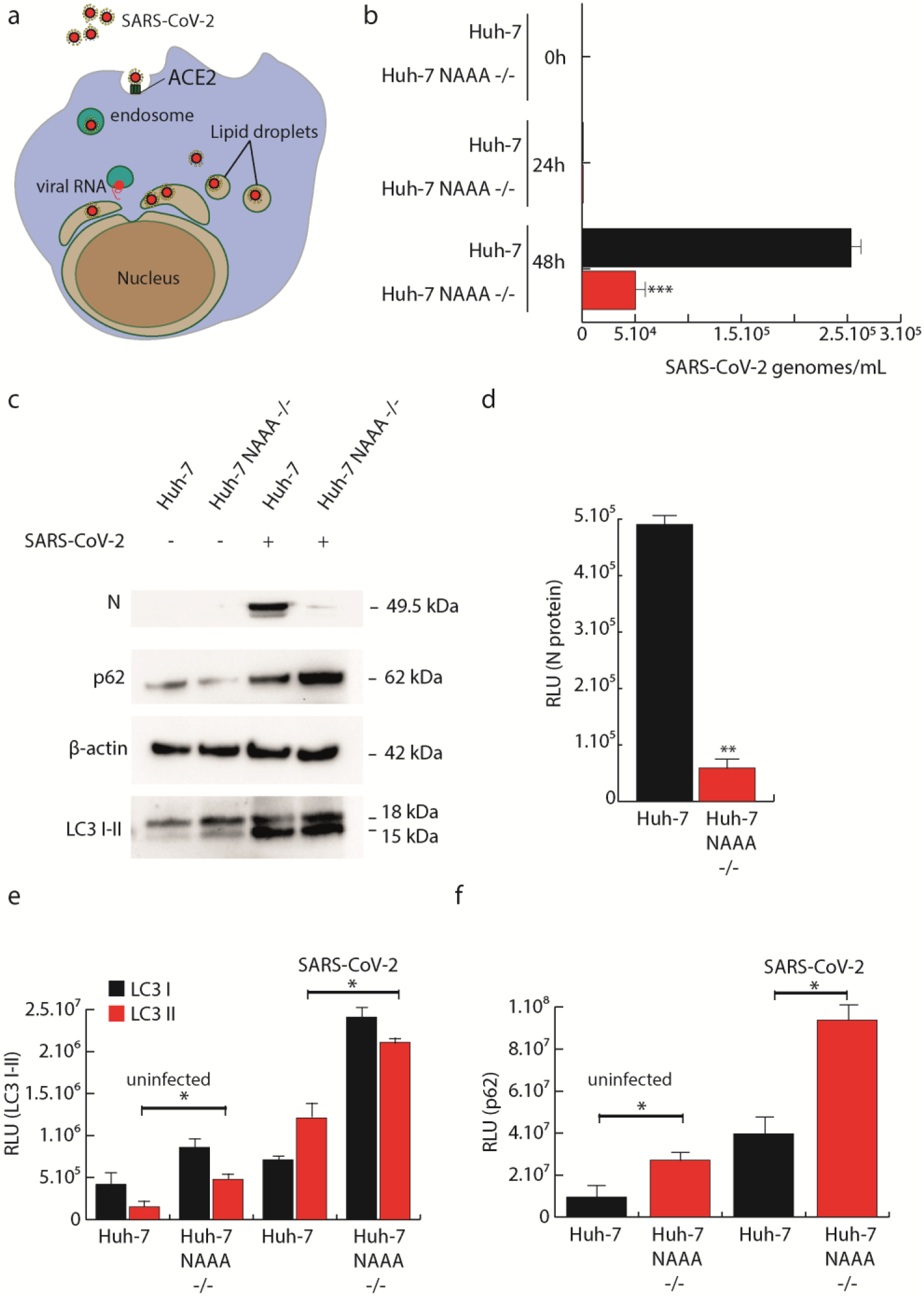
NAAA ablation downregulates SARS-CoV-2 replication. **a.** Schematic illustration of SARS-CoV-2 entry mechanism. Briefly, SARS-CoV-2 complexes with hACE2 receptor. Then, viral S protein is processed either by TMPRSS2 or other serine proteases facilitating endocytosis where S is processed by Cathepsin L in the lysosome. Viral RNA, released from TMPRSS2-mediated entry or endosome release, is replicated as partial and complete genome copies and translated in the endoplasmic reticulum to assemble new SARS-CoV-2 virions. **b.** Time-point qRT-PCR was performed on supernatants taken from Huh-7 and Huh-7 NAAA-/- cells at 0, 24, and 48h after SARS-CoV-2 infection. Statistical analysis was performed with Student’s t-test (* p<0.05, ** p<0.01, *** p<0.001) Data are expressed as mean ± SD, N=4, alpha =0.01. **c.** Western Blot performed on Huh-7 and Huh-7 NAAA-/- before and after SARS-CoV-2 infection. **d-f.** Statistical analysis of SARS-CoV-2 N protein levels (d), autophagic markers LC3 I-II (e), and p62 (f). Statistical analysis was performed using One-Way Anova (* p<0.05, ** p<0.01, *** p<0.001). Data are expressed as mean ± SD, N=4, alpha =0.01.

### NAAA inhibitors exert antiviral activity against multiple variants of SARS-CoV-2

To assess if NAAA inhibitors might reproduce the same antiviral activity as NAAA ablation, we pre-treated Huh-7 cells with ARN077. As illustrated in Figure 6a, the NAAA inhibitor was administered 15 minutes before SARS-CoV-2 infection at the following concentrations: 0.1, 1, and 3 μM. Then, we quantified SARS-CoV-2 genomes in the supernatants of Huh-7 by qRT-PCR (Fig. 6b). Analysis showed a 4-fold decrease of SARS-CoV-2 released virions in the presence of 1 μM ARN077.

**Figure 6.**
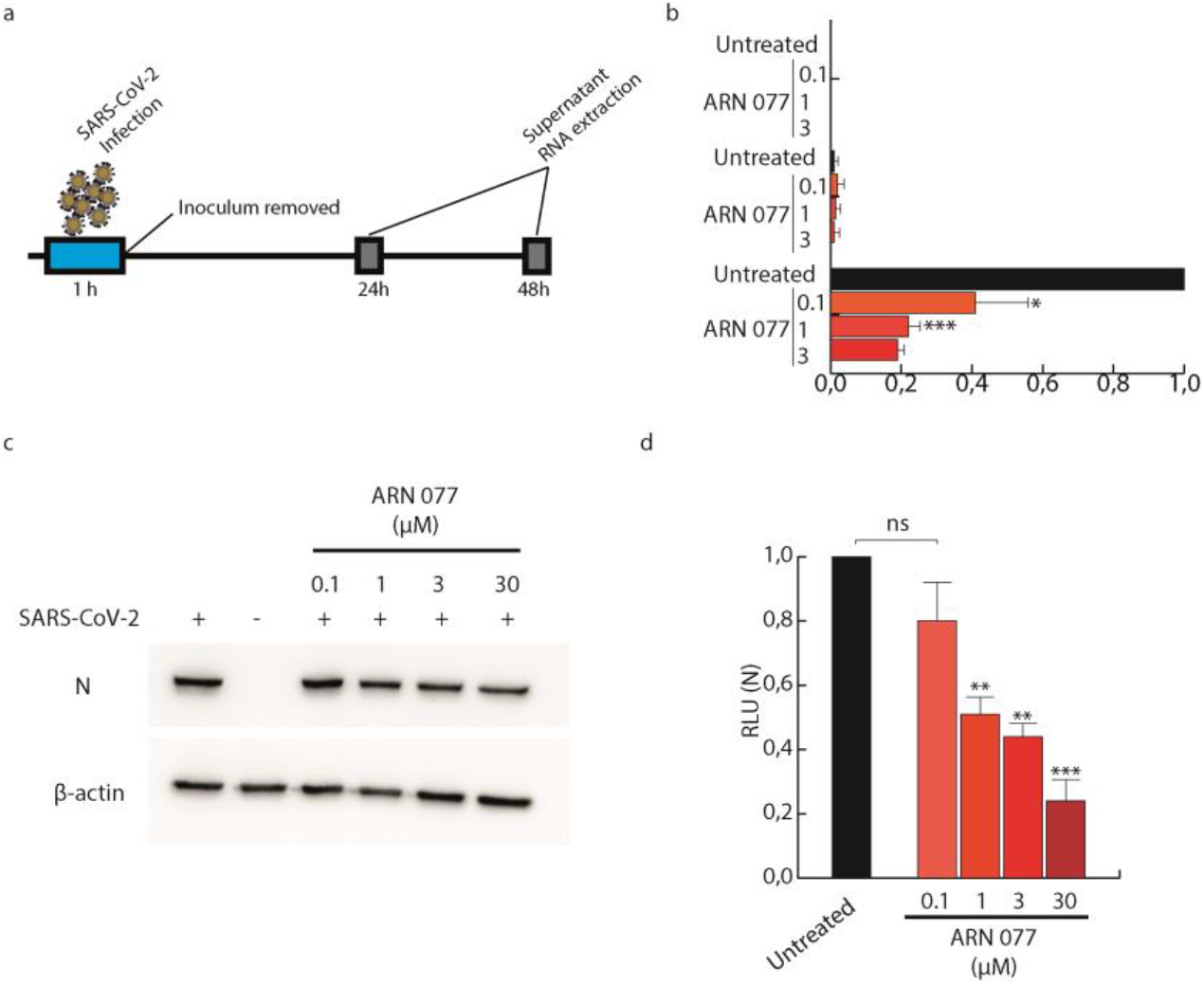
ARN077 decreases SARS-CoV-2 replication. **a.** schematic representation of ARN077 administration. Briefly, we administered ARN077 at various concentrations (0.1, 1, and 3 μM) 15 min prior to SARS CoV-2 infection at 1 MOI. The viral inoculum was subsequently removed and supernatants were collected at 0, 24, and 48h post-infection. **b.** qRT-PCR titration of SARS-CoV-2 virions in the supernatants of Huh-7 cells, treated or not with ARN077. Data were analyzed using One-Way Anova (* p<0.05, ** p<0.01, *** p<0.001) alpha= 0.05, N=6. **c.** Western blot analysis performed on Huh-7 cell lysates treated or not with ARN077 and infected or not with SARS-CoV-2. **d.** N protein content was analyzed using One-Way Anova (* p<0.05, ** p<0.01, *** p<0.001) alpha= 0.05, N=6

Then, to assess whether NAAA inhibition might reduce the replication of SARS-CoV-2 variants, we infected Huh-7 and Huh-7 NAAA-/- cells with two variants of concern, B.1.17 and B.1.617.2, known as *alpha* and *delta* SARS-CoV-2 variants. As previously described, we infected the cells with 1 MOI for 1h. Then the inoculum was removed and replaced with fresh medium, as schematically illustrated in Figure 7a. Total RNA was extracted 48h after infection. As shown in Figure 7b, NAAA ablation decreased replication of both VoC B.1.617.2 and B.1.1.7, as observed previously for the SARS-CoV-2 original strain. Interestingly, Figure 7c shows that SARS-CoV-2 dramatically decreases the PPAR-α receptor (~5 folds) transcripts in WT cells, while infected Huh-7 NAAA-/- cells maintain it at high levels of transcription. The transcription of PPAR-α is known to be increased in response to NAAA inhibition(28); accordingly, it was found up-regulated in the absence of NAAA. Consequently, the peroxisome/virophagy activator PEX-13, the transcription of which is controlled by PPAR-α, drops dramatically in SARS-CoV-2-infected Huh7 WT cells while remaining at high levels in NAAA-/- infected cells.

**Figure 7.**
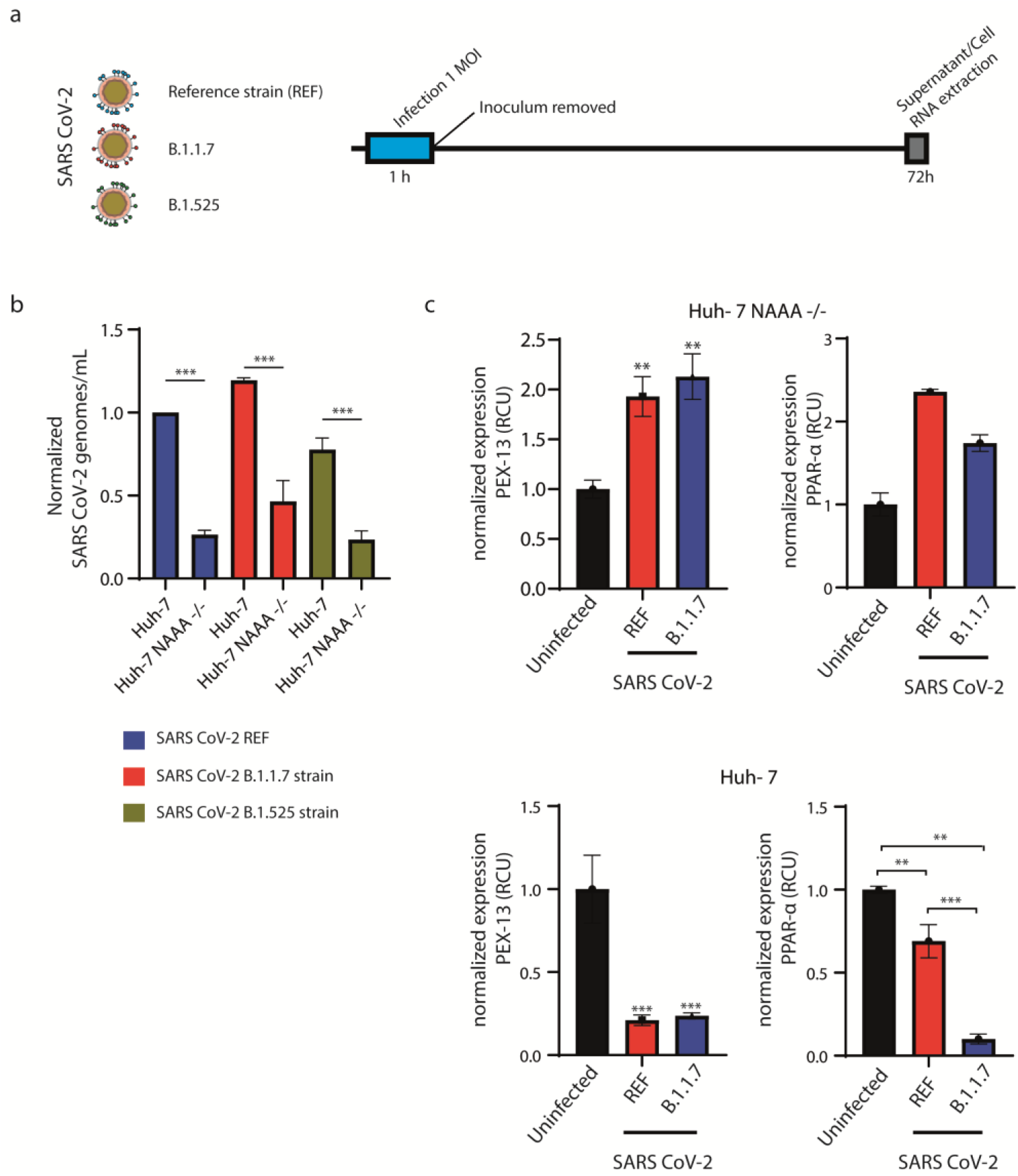
NAAA ablation protects Huh-7 cells from SARS-CoV-2 VoC infection. **a)** Schematic illustration of experimental workflow. Briefly, SARS CoV-2 (Wuhan strain, B.1.1.7 and B.1.617.2) were used to infect Huh-7 and Huh-7 NAAA-/- cells at 1 MOI. Then, supernatants and cell lysates were collected. **b)** qRT-PCR performed on supernatants taken from Huh-7 and Huh-7 NAAA-/- cells 48h after infection with SARS-CoV-2 variants. Data were analyzed with One-Way Anova (* p<0.05, ** p<0.01, *** p<0.001) and are expressed as mean ± SD, N=5, α=0.05. **c)** Normalized expression of PEX-13 and PPAR-α mRNA on Huh-7 and Huh-7 NAAA-/- cells, infected or not with SARS-CoV-2 variants. Data were analyzed with One-Way Anova (* p<0.05, ** p<0.01, *** p<0.001) and are expressed as mean ± SD, N=5, α=0.05.

### ARN077 decreases SARS-CoV-2 infection and inflammation in human PBMC

To confirm the significance of previous data, we tested whether pharmacological NAAA inhibition might decrease infectivity and decrease inflammation on human PBMC taken from healthy donors. To better visualize SARS-CoV-2+ cells, we took advantage of the Prime-Flow technology, an *in situ* hybridization assay that combines branched-DNA technology with the single-cell resolution of flow cytometry (Figure 8a). Briefly, we infected peripheral blood mononuclear cells (PBMC) with SARS-CoV-2, as schematically illustrated in figure 8b. Then we stained cells for CD14+ to identify macrophages and probed them for SARS-CoV-2 genomes and TNF-α. As shown in figure 8c-e, macrophages treated with ARN077 (30 μM) decrease by 70% the amount of SARS-CoV-2 genomes. Remarkably, ARN077 almost abolished SARS-CoV-2-induced TNF-α production, compared with non-treated infected cells.

**Figure 8.**
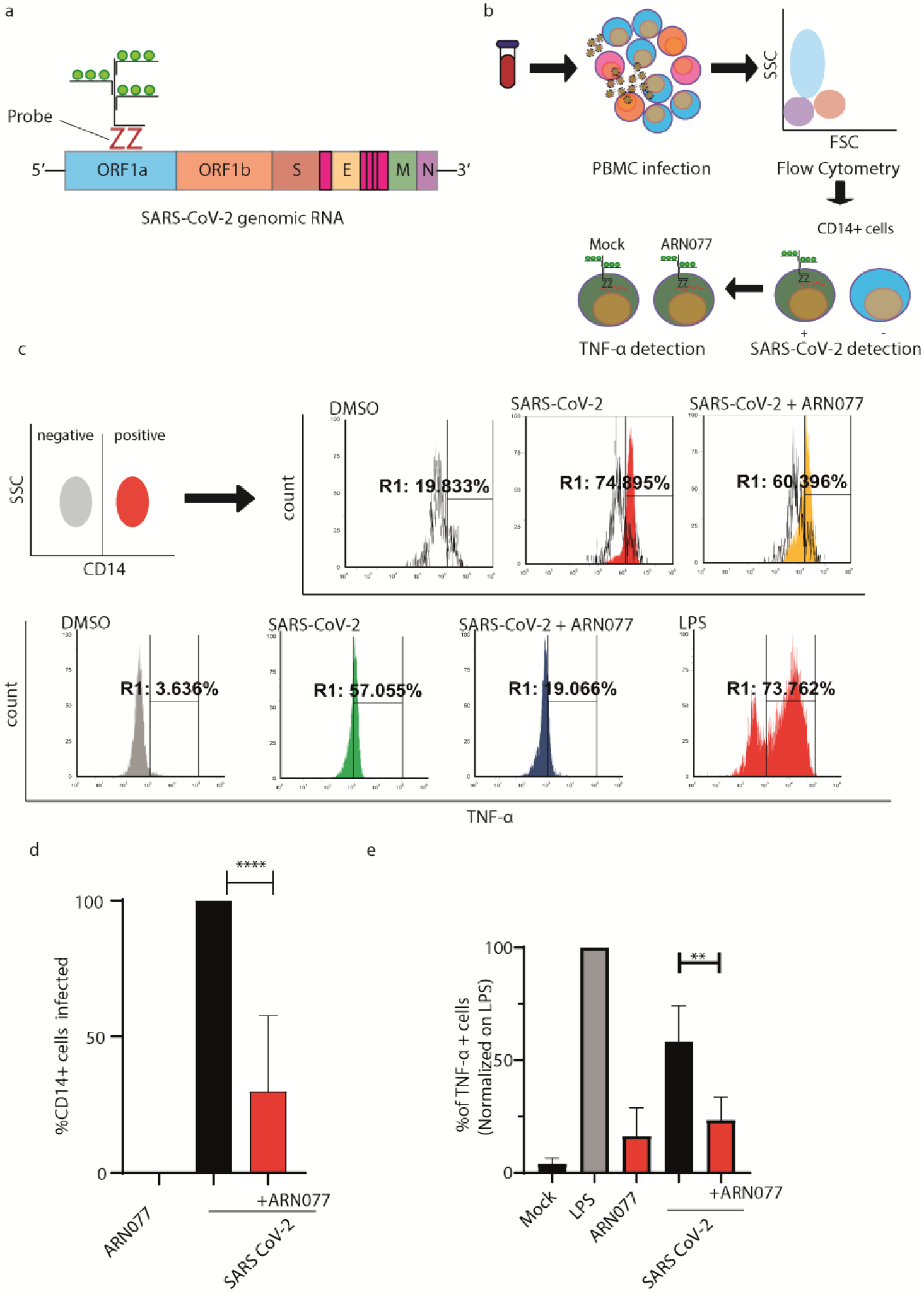
ARN077 reduces SARS-CoV-2 infection and controls inflammation in human PBMCs. **a)** Schematic illustration on the design of SARS-CoV-2 genomic probe, which binds a conserved region of ORF1a. **b**) Experimental workflow of primeflow assay combined with anti-CD14 staining. Briefly, PBMC were infected with SARS-CoV-2, treated or not with ARN077 (30 μM). Then, cells were treated with Brefeldin-α prior fixation and processed by Flow Cytometry. PBMCs were then analyzed for SARS-CoV-2 genomic and TNF-α content. **c**) Representative analysis of PBMC derived macrophages infected or not with SARS-CoV-2. Briefly, CD14+ cells (monocytes and macrophages) were identified by immunostaining. Then, the CD14+ cell population was screened for SARS-CoV-2 intracellular genomes and TNF-α production. DMSO was used as mock-treated control. **d**) Statistical analysis on SARS-CoV-2 genomic content on CD14+ cells, treated or not with ARN077. Data are expressed as mean ± SD and analyzed with One-Way Anova ( **** p<0,0001, n=8, α=0,5) **e**) Statistical analysis performed on % of TNF-α+ cells, treated or not with ARN077 and infected or not with SARS-CoV-2. Data were normalized on cells treated with Lipopolysaccharide (LPS). Data are expressed as mean ± SD and analyzed with One-Way Anova (** p<0,01, n=8, α=0,5).

### NAAA ablation promotes autophagy and increases vesicular pH

Prompted by the evidence that autophagy remains active in NAAA-/- cells during ZIKV and SARS-CoV-2 infection, we decided to dissect the PPAR-α pathway, which is activated by intracellularly accumulated PEA. Since PEA is mainly degraded by NAAA, the absence of the latter causes PEA accumulation; in line with this fact, we observed a 2-fold increase in PPAR-α expression in NAAA-/- cells, as measured by qRT-PCR (Fig. 10a). This transcription factor, once activated by PEA, enhances its transcription by a positive feedback loop. Interestingly, PPAR-α also enhances transcription of PEX-13, a potent selective inductor of viral autophagy (virophagy)(29). Instead, the levels of expression of PEX-14 and PEX-19, which are required for general autophagy, were found unchanged.

**Figure 10.**
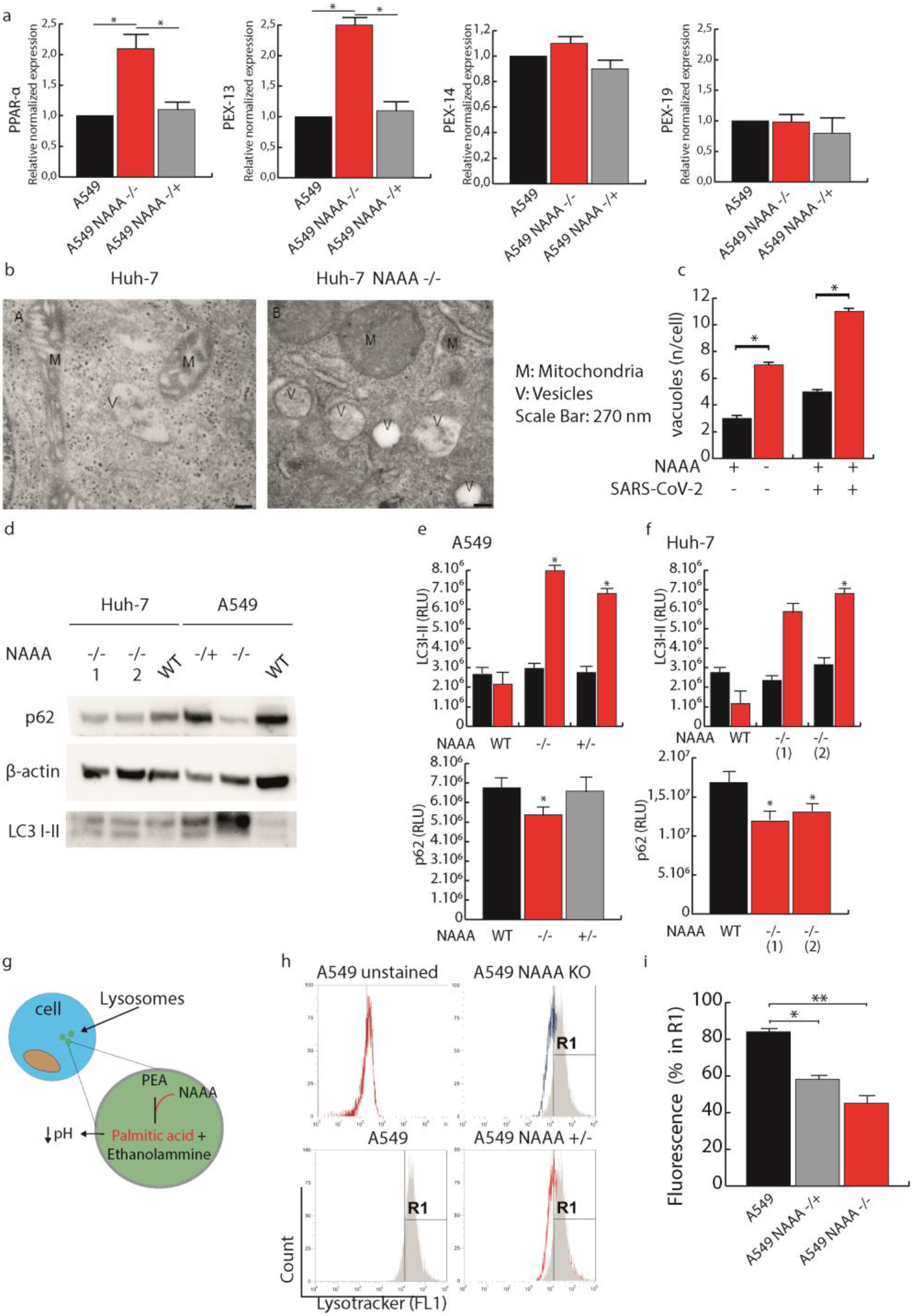
NAAA suppression promotes autophagy through PPAR-α activation. **a.** Quantification of PPAR-α and peroxine (PEX-13, −14, −19) expression in A549 NAAA-/-, +/- and WT cells. Data are expressed as mean ±SD, N=4, and analyzed with One-way Anova (* p<0.05). **b.** Electron microscopy analysis of intracellular vesicles in Huh-7 NAAA-/- and their WT counterparts. **c.** Statistical analysis of intracellular vacuole content described in (b). Data are expressed as mean ±SD, N=4, and mean±S.E.M., N=30 and analyzed with One-way Anova (* p<0.05). **d.** Western blot analysis of autophagy markers p62 and LC3 I-II in A549/Huh-7 NAAA -/-, +/- and WT cell lysates. **e-f.** Statistical analysis of p62 and LC3-II levels shown in d. Data are expressed as mean ±SD, N=4, and analyzed with One-way Anova (* p<0.05). **g.** Schematic illustration of intra-vesicular degradation of PEA. NAAA hydrolyzes PEA into palmitic acid and ethanolamine, contributing to the acidification of lysosomes. **h.** Representative flow cytometry acquisition of A549, A549 NAAA-/- and NAAA+/- cells stained with lysotracker dye. **i.** Statistical analysis of flow cytometry acquisition of A549, A549 NAAA-/- and NAAA+/- cells stained with lysotracker dye. Data are expressed as mean ± SD, analyzed with One-way Anova (* p<0.05, ** p<0.01, α=0.1).

As shown in Figures 10b and c, electron microscopy analysis revealed that Huh-7 NAAA-/- cells possess 5 times more intracellular vesicles than their WT counterparts. Moreover, we probed both A549 and Huh-7 for LC3-II and p62 protein content. As shown in Figure 10 d-f, we found that both NAAA-/- cells (Huh-7 and A549) increase the levels of LC3-II, while the cargo protein p62 decreases in both cell lines. Based on autophagy guidelines^r(30)^, this condition indicates an active autophagic flux of both A549 and Huh-7 NAAA-/- cells.

Finally, to evaluate vesicular pH variations, we performed A549 NAAA-/- live cell staining with lysotracker dye, which increases fluorescence when pH decreases. Since the absence of NAAA is likely to reduce palmitic acid within vesicles, this expected to produce a lower acidic environment, as shown in Figure 10 g. Lysotracker assay revealed that both A549 +/- and -/- decrease the vesicular acidity (Fig. 10 h-i).

### NAAA inhibition maintains active autophagy during SARS CoV-2 replication

To further confirm the status of autophagy during SARS-CoV-2 infection and autophagy induction when NAAA activity is impaired, we analyzed Huh-7 WT and Huh-7 NAAA-/- cells. This allows to detect LC3 positive vacuoles and LC3 particles within vacuoles and cytosol, together with SARS-CoV-2 virions by transmission electron microscopy (TEM). As reported in representative Figure 11a, SARS-CoV-2 virions (black arrows) are widely dispersed within the cytoplasm of Huh-7 cells. In contrast virions are found within vacuoles in Huh-7 NAAA-/- cells. Remarkably, virions (red arrows of Figure 11b) are widely dispersed within the cytosol of Huh-7 infected cells, while in NAAA-/- cells virions are compartmentalized within LC3 positive vacuoles. LC3 immuno-electron microscopy (Figure 11b), shows that Huh-7 NAAA-/- cells own an increased amount of LC3 positive vacuoles which is significantly higher compared with WT Huh-7 cells as counted in graph of Figure 11c. The amount of LC3 positive vacuoles is reduced during SARS-CoV-2 infection in Huh-7 NAAA WT cells, however this reduction is prevented within Huh-7 NAAA-/- cells. This is measured in Figure 11c, where it is evident that NAAA deletion drastically increases the number of LC3 positive vacuoles. This occurs in both groups of NAAA-/- cells, independently from the presence of SARS-CoV-2. In detail, as measured in the graphs of Figure 11c, LC3 positive vacuoles increase roughly three-fold within NAAA-/- cells compared with WT.

**Figure 11.**
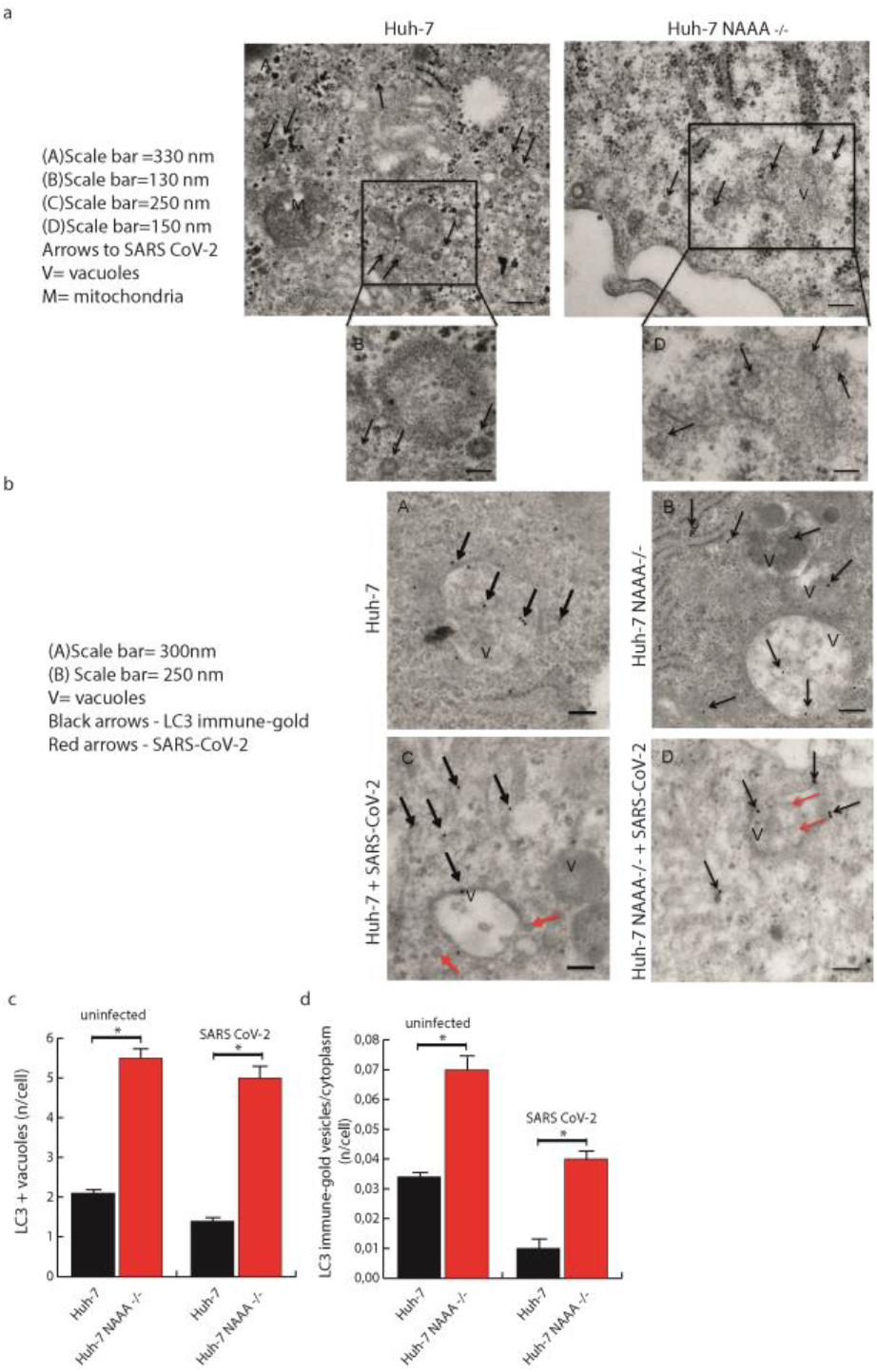
Transmission electron microscopy of Huh-7 and Huh-7 NAAA-/- infected with SARS-CoV-2. **a.** Representative micrographs showing Huh-7 WT and Huh-7 NAAA-/- cells infected with SARS-CoV-2 using plain transmission electron-microscopy (TEM) **b.** Representative micrographs showing of Huh-7 and Huh-7 NAAA-/- infected or not with SARS-CoV-2 and probed with LC3+-immune gold (immune-electron microscopy) **c.** Counts of LC3 positive vacuoles per cell, performed within Huh-7 WT and Huh-7 NAAA-/-, infected or not with SARS-CoV-2. **d.** Counts of the ratio between stoichiometry amount of LC3 particles within vacuoles/LC3 particles within cytosol within Huh-7 WT and Huh-7 NAAA-/-, infected or not with SARS-CoV-2. Data are expressed as the mean ± S.E.M., N=30 (* p<0.05). Statistical analysis was performed using ANOVA. Data are expressed as mean ± SD, N=3, α=0.1. (* p<0.05)

To further analyze the autophagy status within NAAA-/- cells during SARS-CoV-2 infection, we calculated the ratio between LC3 immuno-gold particles within vacuoles and within cytosol (Figure 11d). In all cells SARS-CoV-2 infection decreases the compartmentalization of LC3 within vacuoles compared with cytosol (Figure 11d). However, within NAAA-/- cells the compartmentalization of LC3 (vacuoles vs cytosol) is much higher in NAAA-/- Huh-7 infected cells compared with WT Huh-7 infected cells (Figure 11d). This indicates that when NAAA expression is inhibited a higher amount of LC3 is compartmentalized within vacuoles, which contrast effectively the decreased compartmentalization induced by SARS-CoV2. Thus, during SARS-CoV2 infection a general loss occurs for vacuolar compartmentalization of LC3, however such a loss is counteracted effectively by deleting NAAA. In these latter condition the ratio of LC3 within vacuoles and LC3 within cytosol is rescued to a level similar to non-infected Huh-7 cells (Figure 11d). Altogether data from TEM and ultrastructural stoichiometry indicate that: (i) deletion of NAAA from Huh-7 cells is able to increase autophagy vacuoles, (ii) deletion of NAAA counteracts the loss of LC3 compartmentalization, which is produced by SARS-CoV2 infection.

The loss of LC3 compartmentalization, which is induced by SARS-CoV-2 infection is much more marked when compared with the slight decrease in LC3 immuno-positive vacuoles, this indicates that the decrease in LC3 particles mostly occurs in the vacuolar compartment during SARS-CoV-2 infection

### NAAA inhibition promotes β-oxidation through PPAR-α activation

PPAR-α, once activated by PEA, increases its transcription by positive-feedback loop(31). In turn, PPAR-α is known to enhance the transcription of several mediators of β-oxidation. Among them, CPT1a, HADHa, LCAD, and UCP2 modulate mitochondrial β-oxidation, while ACOX increases peroxisomal β-oxidation(32). The increased intracellular content of LC3b, previously described for NAAA-depleted cells, might activate selective autophagy of lipid droplets (LD)(33). Figure 12a schematically shows how PPAR-α activates various pathways involved in LD dismantling.

**Figure 12.**
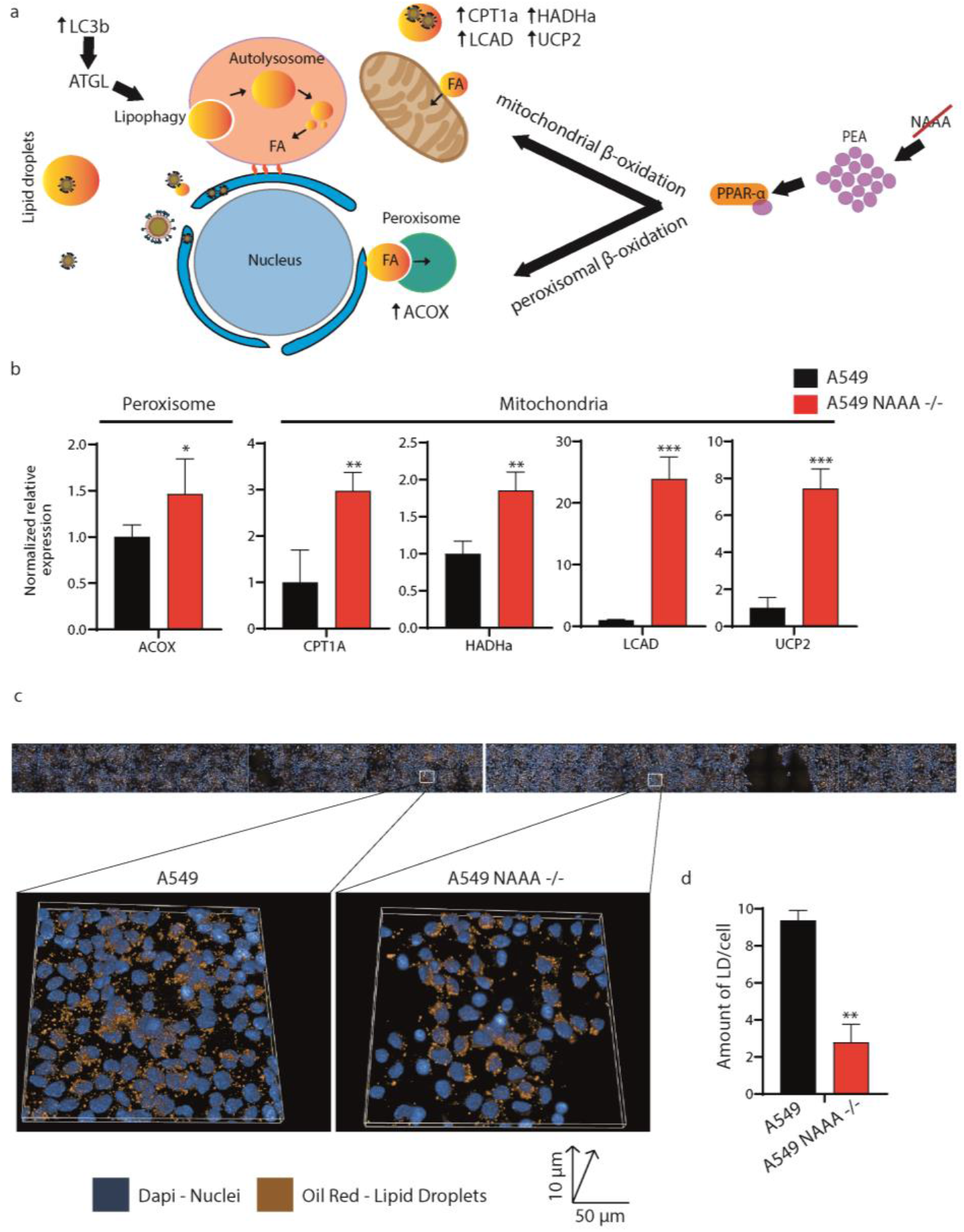
NAAA ablation dismantles intracellular LDs. **a.** Schematic illustration of PPAR-α induction of β-oxidation. Black arrows indicate genes that are directly regulated by PPAR-α. **b.** qRT-PCR performed on A549 and A549 NAAA-/- cells, normalized on β-actin. Data are expressed as mean ± SD and analyzed by Student’s *t*-test (N=3, * p<0.1, ** p<0.01, *** p<0.001). **c.** High content screening performed on A549 and A549 NAAA -/- stained with OilRed (orange). Nuclei were stained with DAPI, a section of 10 μ was analyzed in 40 fields per cell type. Scale bar: 50 μ. **d.** Statistical analysis performed on LD content/cell in both A549 and A549 NAAA-/- cells. Data are expressed as ± SD, normalized on the number of nuclei, and analyzed by Student’s *t*-test (N=3, * p<0.1, ** p<0.01, *** p<0.001)

To probe the hypothesis that NAAA inhibition might dismantle LD needed for SARS-CoV-2 and ZIKV replication, we measured transcription of PPAR-α by qRT-PCR. As shown in Figure 12b, A549 NAAA-/- cells exhibit increased transcription of genes involved in both mitochondrial and peroxisomal β-oxidation; particularly, expression of LCAD and UCP2 were found 20 and 5 times higher, respectively, in NAAA-/- than in WT cells. CPT1A and HADHa showed at least double the expression than in WT counterparts, while ACOX was moderately but still significantly augmented. To confirm increased β-oxidation at the lipid level, we stained LD in both A549 and A549 NAAA-/- with OilRed, expected to selectively mark LD intracellularly. Next, we took advantage of high content confocal screening to count LD intracellular content three-dimensionally and objectively count them. As shown in Figure 12c, LD content drops abruptly ~5 times in NAAA-/- cells, compared to their WT counterparts.

## Discussion

The spread of SARS-CoV-2 worldwide represents a threat to global public health. By October 4^th^, 2021, SARS-CoV-2 had infected 234,809,103 people and had caused 4,800,375 deaths around the world (WHO, 2021). In this context, we face the absence of specific antiviral drugs and the urgency of novel and effective treatments against emerging viruses. More generally, given the hastiness with which drugs are required during sudden pandemics, it would be highly desirable to repurpose drugs already used for clinical treatments which do not require further toxicity studies.

In the present work, we tested an antiviral drug candidate, which might exert two distinctive, seminal functions: blocking viral replication and, at the same time, avoid excessive inflammatory response.. For this reason, we investigated the antiviral effectiveness of NAAA inhibitors, well-established anti-inflammatory, and antinociceptive drugs(9).

Here, we show that SARS-CoV-2 and ZIKV reduce their replication by ~5 folds in different cell lines, when the NAAA enzyme is inhibited using pharmacological or gene knock-out techniques. Once the NAAA enzyme is blocked, ZIKV and SARS CoV-2 decrease their replication and, consequently, also virion release drops abruptly by ~5 folds for both viruses. Surprisingly, we also found that NAAA inhibition maintains autophagy active during infection, whereas both SARS-CoV-2 and ZIKV viruses would hijack this pathway to boost their replication(25, 26, 34). A possible explanation for such an effect is that, in the context of NAAA inhibition, neither virus replicates to a level sufficient to counteract autophagic activation. Therefore, active autophagy contributes to viral clearance. This may appears unexpected at first glance, since both SARS-CoV-2 and ZIKV viruses would hijack early autophagy structures to boost their replication. It is possible that the stimulus induced by NAAA inhibition to address autophagy-related membrane web towards lysosomal degradation counteract the slowed autophagy flux in the course of viral infection, and it may explain the suppression of viral replication even when occurring within early autophagy structures.

Autophagy-related proteins can be targeted by different RNA viruses(35, 36). While coronaviruses and flaviviruses use autophagic induction to produce double-membrane vesicles in which to replicate, the latter is not required for COXB5 replication that exploits other autophagy-independent intracellular membranes(37). Accordingly, we show that NAAA inhibitors do not reduce replication of COXB5, possibly because this virus flexibly uses autophagic membranes and others of different origins, in agreement with the previous findings(37).

Therefore, NAAA ablation results in antiviral activity by two distinct routes. First, the accumulation of NAAA substrate, PEA, activates PPAR-α, a transcription factor that once activated generates a cascade of events that lead to the disruption of fatty acid droplets(16), thereby bringing about LD destruction through β-oxidation. These, in contrast, are essential for the replication of flaviviruses (17) and, as recently suggested, also for SARS-CoV-2(18). Secondly, NAAA inhibition maintains autophagic flux active as a second line of defense to counteract viral assembly in infected cells. Our data point out that NAAA inhibition induces virophagy through positive regulation of transcription of PEX13, one of its major activators(29).

Recent evidence demonstrates that SARS-CoV-2 triggers reprogramming of lipid metabolism in monocytes and other cells, thus accumulating LDs to favor virus replication(18). Moreover, the inhibition of LD biogenesis modulates viral replication and pro-inflammatory mediator production(18). As expected, we show that NAAA inhibition dismantles LDs, probably diminishing the possible sites of viral assembly.

It has long been known that macrophage M2 polarization is dependent on fatty acid oxidation and that inhibition of cell-intrinsic lysosomal lipolysis suppresses M2 activation during infection with a parasitic helminth(38). The SARS-CoV-2 infection has been recently described to trigger pathological profibrotic macrophage polarization(39), perhaps by triggering reprogramming of lipid metabolism and accumulation of LDs. Further, it has been shown that perturbation of LD biogenesis influences viral replication and pro-inflammatory mediator production(18). As expected, we show that NAAA inhibition leads to reduce SARS-CoV-2 genomic content in the very cells where SARS-CoV-2 and ZIKV replicate, macrophages, which are also key players in pathogenesis. NAAA might be a key player in the therapeutical control of macrophage polarization. Ongoing studies are aimed at questioning whether monocyte expression of CD163, a macrophage marker of pro-fibrotic activity, might be influenced by NAAA inhibitors in SARS-CoV-2 infected monocytes.

Our findings support the notion that host lipid metabolism and LDs are required for ZIKV and SARS-CoV-2 replication and suggest a novel potential strategy to interfere with viral replication in infected cells by blocking NAAA is a potential novel strategy to counteract viral replication in cells infected by at least ZIKV and SARS-CoV-2.

Together, these results highlight that a well-recognized anti-inflammatory agent can be also used as an antiviral drug for the first time. Our data show that peripheral blood-derived human monocytes infected with SARS-CoV-2 and treated with ARN077 exhibit decreased viral genome content together with reduced production of TNF-α, compared to ARN077 untreated counterparts. This finding supports the hypothesis that NAAA inhibition might be beneficial to treat COVID19 patients and, hopefully, prevent aberrant macrophage activation during acute respiratory distress syndrome. While NAAA was never studied as an antiviral target until now, its metabolite PEA has been used in several placebo-controlled double-blind clinical trials on influenza and the common cold. Promising results led to the clinical use of PEA under the brand name Impulsin in former Czechoslovakia (4). As PEA plays a fundamental role as an anti-inflammatory lipid-modulating precursor, its activity is counteracted by NAAA, which is abundant in macrophages and immune cells(40), the very cells responsible for cytokine storm in severe COVID19. For this reason, while the antiviral activity of PEA might not be durable, NAAA inhibition might have a longer-lasting efficacy and more potent anti-inflammatory/antiviral activity, especially when administered in combination with PEA.

In summary, this work unveils a critical interaction between NAAA enzyme and viral replication, proposing a new mechanism that ZIKV and SARS-CoV-2 use to hijack the host innate immune defenses. Moreover, we describe a new antiviral strategy based on the suppression of NAAA and of the cascade of events leading to blockage of replication of ZIKV and SARS-CoV-2, while maintaining an anti-inflammatory environment. This dual activity might be exploited in clinical practice to both reduce SARS-CoV-2 replication and the cytokine storm it causes, which is still one of the major pathogenic events that lead to COVID-19 lethality.

## Materials and methods

### Cell Culture, transfection, and treatments

A549 and Huh-7 cells were purchased from American Type Culture Collection (Manassas, VA 20110, USA), and cultured at 37 °C in 5% CO2 in Dulbecco-Modified Eagle’s Medium (DMEM) supplemented with 10% fetal bovine serum (FBS), 100 U/mL penicillin and 100 μg/mL streptomycin. Cells were seeded in 96-well plates and transfected 24h after with pMXs GFP-LC3-RFP (#117413, Addgene, MA) or with Polyinosinic-polycytidylic acid sodium salt, Poly (I:C) (0.5 ng/μL), using Lipofectamine LTX and PLUS reagent (Thermo Fisher Scientific, Waltham, MA 02451, USA) according to the manufacturer’s instructions. NAAA inhibitor ARN077 was kindly provided by Prof. Daniele Piomelli. Cells were tested for mycoplasma contamination as previously described(41)

### CRISPR/Cas9 design and transfection

A549 and Huh-7 cells were transfected with CRISPR/Cas9 RNP (IDT, Coralville, Iowa) by nucleoporation (NEON Electroporation System, Thermo Fisher, Massachusetts, USA) using the following parameters: 1200 V, 30 ms, 2 pulses using the sgRNA guide: AGTGGGTGCACGTGTTAATC. To isolate single-cell clones, cells were seeded in 96-well performing a single-cell limiting dilution protocol as previously described(22, 23). Edited clones were selected by sequencing analysis of PCR products, using the following primers: F 5’-GAGGCTGCAGATTGAGTGAC-3’, R 5’-TGGAGGTTTCTAGGCAAGCA-3’.

### Infections

Cells were infected using Zika virus (strain MP1751, Uganda), SARS-CoV-2 (VR PV10734; B.1.1.7 strain; B.1525 strain and COXB5 (isolated at Cisanello hospital, Pisa, Italy). A549 and Huh-7 cells and NAAA-/- cells were infected with 1 M.O.I of Zika or SARS-CoV2. A549 and A549 NAAA-/- cells were infected with COXB5 at 0.5 MOI.

### Flow cytometry

A549 and A549 NAAA-/- cells (50.000 cell/well) were seeded in 24-well plate and incubated overnight. Then, cells were stained with LysoTracker™ Green DND-26 (L7526, Thermo Scientific) following the manufacturer’s instruction and analyzed by flow cytometry using ATTUNE NXT Flow Cytometer (Thermo Fisher Scientific, Waltham, MA, USA).

### Western blot analysis

A549 and Huh-7 cells were lysed with RIPA lysis buffer (Millipore, Massachusetts, USA). membranes were incubated at 4 °C overnight with the following antibodies: anti-NAAA (1:1000, Abnova, Taipei, Taiwan), Anti-ZIKV E (1:1000, Genetex; Irvine, US) anti-SARS-CoV-2 N (1:1000, MA14AP1502, Sino Biological, Beijing, China) anti-P62 (1:1000, ab56416, Abcam, Cambridge, UK), anti-LC3 I-II (1:1000, L7543, Sigma-Aldrich, St. Louis, MO 63103, USA), anti-ATG5 (1:1000, Anti-APG5L/ATG5 antibody, ab228668, Abcam, Cambridge, UK) and anti-β-actin (1:1000, A2066 Sigma-Aldrich, St. Louis, MO 63103, USA). Blots were acquired and analyzed by using Chemidoc XRS system (BioRad, California, USA).

### RNA extraction and qRT-PCR

Total RNA was extracted with QIAzol Lysis Reagent (QIAGEN, Hilden, Germany) according to the manufacturer’s instructions. Total RNA (200 ng) was reverse-transcribed in cDNA and amplified by using QuantiNova SYBR Green RT-PCR kit (QIAGEN®, Hilden, Germany). Viral genomes were extracted from the supernatants at different time points post-infection using Takara MiniBEST Viral RNA/DNA (Takara Bio, Kyoto, Japan) according to the manufacturer’s instructions. Then RNA was reverse-transcribed and amplified using One Step PrimeScriptTM III RT-qPCR Mix kit (Takara Bio, Kyoto, Japan). Primers and probes: ZIKV: F:5’-TGAGATCAACCACTGCAAGY-3’, R: 5’-GCCTTATCTCCATTCCATACCA-3’, Probe 5’- FAM-ATCGAGGAATGGTGCTGCAGGGA-BHQ1-3. SARS-CoV-2: F: 5’-TCACCTATTTTAGCATGGCCTCT -3, R: 5’-CGTAGTGCAACAGGACTAAGC-3, Probe: 5’-/56-FAM/TGCTTGTGCCCATGCTGC-3’. NAAA: F:5’-AAGACTCCAGAGGCCACATTTACCATGGTC-3’; R:5’CATCAGCAATAAGGGGAGTCTTGGCCAACT-3. PPAR-α: F: 5’-CTATCATTTGCTGTGGAGATCG-3’, R: 5’-AAGATATCGTCCGGGTGGTT-3’ Pex13: F:5’-GGGCCCCATTTTCCAATCCTG-3’, R:5’-TACACGGAGGCGGTTGTAGC-3’ Pex14: F:5’-GCCACGGCAGTGAAGTTTCTA-3’, R:5’GCTGGAAGGCCATATCAATCT-3’ Pex19: F:5’-GATCACAGAAAAGTATCCAGAATGGTT-3’, R:5’-CGAGCCTTTTGAGTGGTTTCA-3’ HADHA: F:5’-GCTAGACCGAGGACAGCAAC -3’, R:5’-CCTGCTTGAGACCAACTGCT-3’ CPT1a: F:5’-TGAGCGACTGGTGGGAGGAG -3’, R: 5’-GAGCCAGACCTTGAAGTAGCG-3’ UCP2: F: 5’ CACCAAGGGCTCTGAGCATG -3’, R: 5’-TCTACAGGGGAGGCGATGAC-3’ ACOX1: F: 5’ TCCTGCCCACCTTGCTICAC -3’, R: 5’-TTGGGGCCGATGTCACCAAC-3’ LCAD: F: 5’-TGAGCGACTGGTGGGAGGAG-3, R: 5’-CACTGTCTGTAGGTGAGCAACTG - 3’ β-actin: F: 5’AGGAGAAGCTGTGCTACGTC-3’, R: 5’-AGACAGCACTGTGTTGGGGTA-3’.

### Immunostaining and High-content confocal imaging

A549 or Huh-7 cells were seeded in 96-CellCarrierUltra plates (Perkin Elmer, Hamburg, Germany), then infected or treated as described above. Cells were fixed using 3.7% formaldehyde and stained with the following primary antibodies: anti-NAAA (1:300, Abnova, Taipei, Taiwan), ZIKA: Anti-Flavivirus NS1 antibody (ab214337), Anti-SARS-CoV-2 spike protein (Sino Biological, Beijing, China, 1:200). LDs were stained using OIL Red R (1:5000 diluted in water) for 30 min (Sigma-Aldrich, St. Louis, USA). Nuclei were stained with DAPI (1μg/mL). Images were acquired using Operetta CLS high-content imaging device (PerkinElmer, Hamburg, Germany) and analyzed with Harmony 4.6 software (PerkinElmer Hamburg, Germany). To investigate the co-localization of ZIKV (NS1) and NAAA we used the following building blocks: Find Nuclei > Find Cytoplasm > Calculate intensity properties (ZikvNS1-Alexa 488) > Select population: Infected cells > Find Spot NS1+ > Find Spot NAAA+ > Calculate position properties (% overlap NS1+/NAAA+). Similarly, co-localization with autophagosomes/autolysosomes was performed with the following building blocks: find nuclei > find cytoplasm > Calculate intensity properties (GFP, RFP, NAAA) > find spots (GFP+RFP, GFP-/RFP+) > calculate position properties (%overlap GFP+/RFP+ and NAAA, GFP-/RFP+ and NAAA.) Twenty-five fields were analyzed per well using 63× water objective.

### Transmission electron microscopy (TEM)

Huh-7 cells were infected or treated as described above. Then, cells were fixed with 2.0% paraformaldehyde/0.1% glutaraldehyde, both dissolved in 0.1 M PBS pH 7.4 for 90 min at 4 °C. Cells were gently scraped from the plate, collected into vials, and centrifuged at 10,000 rpm for 10 min. cells were then resuspended in PBS and post-fixed in 1% osmium tetroxide (OsO4) for 1 h at 4 °C. Then, the post-fixed pellet was dehydrated in a gradient of ethanol solutions 50%, 70%, 90%, and 95%, each for 5 min, to reach 100% ethanol for 60 min. Finally, the pellet was embedded in epon-araldite for 72 h, at 60°C. The embedded cell samples were sectioned at the ultramicrotome (Leica Microsystems) and ultrathin sections underwent conventional electron microscopy or immune-electron microscopy. Samples were observed at Jeol JEM SX100 TEM (JEOL, Tokyo, Japan) at an acceleration voltage of 80 kV.

### Human peripheral blood mononuclear isolation and infection

Clinical study was approved by local ethics committee (protocol number 19204) in accordance with institutional guidelines; participants were aware of the nature of the study and signed written informed consent), PBMCs derived from healthy donors (age 30-60, no chronic comorbidities, absence of medications) were isolated by gradient centrifugation using Lympholite-H Cell Separation Media. PBMCs (2 × 10^7^) were collected into polypropylene conical tube for Flow Cytometry and incubated for 24 h at 37°C, 5% CO2. PBMCs were activated with LPS 1 ng/mL for 4 h in the presence with Brefeldin-A. PBMCs were pre-treated or not with ARN726 for 15 min and infected with SARS CoV-2 0,5 MOI for 48 hours. Then, cells were collected and analyzed with Prime Flow RNA assay according to the manufacture’s instruction. RNA probe to detect SARS-CoV-2 were designed to bind the ORF1a genomic region. Cells were stained against Anti-CD14 PE-Cyanine-7 to detected monocyte population, and anti-TNFα-PE.

### Post-embedding immune-electron microscopy and data analysis

The localization of specific proteins at the ultrastructural level using gold-conjugated antibodies is a valuable technique based on our previous studies(42, 43). Briefly, we carried out sample preparation to preserve antigenicity besides the maintenance of a good ultrastructure morphology. Ultrathin sections were collected on a nickel-coated grid, processed for LC3 detection using a primary antibody (Rabbit anti-LC3, Abcam, ab128025, AB_11143008). Ultrathin sections were incubated on droplets of aqueous sodium metaperiodate (NaIO4) for 30 min, at 22 °C to remove OsO4. After washing in PBS, grids were incubated in drops of blocking solution (10% goat serum and 0.2% saponin in PBS) for 20 min at 22 °C. Then, in a humidified chamber, the grids were placed on drops containing the primary antibody (diluted 1:50 in ice-cold solution in PBS containing 1% goat serum and 0.2% saponin) overnight at 4 °C. After washing in cold PBS, ultrathin sections were incubated in the gold-conjugated secondary antibodies (20 nm gold particles, EM Goat anti-Rabbit IgG, gold conjugated antibody BBInternational EM.GAR 20 AB_1769136), diluted 1:20 in blocking buffer (1% goat serum and 0.2% saponin in PBS) for 1 h at 22 °C. Counts of LC3 immune-gold particles (20 nm) were performed at TEM at the minimal magnification (8000X) in which both the immune-gold particles and the cell organelles can be identified(44, 45). Grids were analyzed to count immune-gold particles within 30 cells for each experimental group. Ultrastructural morphometry assessments of vacuoles and measurement of immunogold particles were carried out as previously described^40^. Values were expressed as follows: (i) total number per cell of LC3 immune-gold particles, (ii) number of unstained vacuoles per cell, (v) number of LC3-positive vacuoles per cell, and (vi) ratio of the number of LC3 immune-gold particles within vacuoles out of the number of cytoplasmic LC3 immune-gold. Data were expressed as the mean value±S.E.M. and were analyzed with One-way Anova. Null Hypothesis H0 was rejected when p<0.05.

## Acknowledgments

This work was supported by *“SENSOR, nuovi sensori Real-Time per la determinazione di contaminazioni chimiche e microbiologiche in matrici ambientali e biomedicali”*, Progetto co-finanziato dal POR FESR Toscana 2014-2020; “*Addressing viral neuropathogenesis: Unraveling the molecular and cellular pathways of viral replication and host cell response and paving the way for the development novel host-targeted, broad spectrum, antiviral agents*”, bando PRIN: Progetti di ricerca di rilevante interesse nazionale, Bando 2017, Prot. 2017KM79NN; “*I-GENE, In-vivo Gene Editing by Nanotransducers*”, European call identifier H2020-FETOPEN-2018-2020, Proposal ID 862714. University of Pisa Grant: PRA_2020_37; We acknowledge CISUP—Centre for Instrumentation Sharing—University of Pisa for the use of Operetta CLS imaging facility.

## Contributions

M.L., D.P., G.F and M.P. conceived and designed the experiments; V.L.R., R.A., E.I., C.F., E.C., L. B., R.F., G.L., P.G.S., P.L., S.M., A.M and PQ performed and analyzed the experiments; F.F., M.L., G.F., D.P. and M.P. supervised analyses and provided reagents and resources; M.P. provided funding; M. L., G.F., and M.P. wrote the manuscript with input from all other authors. We declare that we have no conflict of interest.

